# A conserved role for SFPQ in repression of pathogenic cryptic last exons

**DOI:** 10.1101/2020.03.18.996827

**Authors:** Patricia M. Gordon, Fursham Hamid, Eugene V. Makeyev, Corinne Houart

## Abstract

The RNA-binding protein SFPQ plays an important role in neuronal development and has been associated with several neurodegenerative disorders, including ALS, FTLD, and Alzheimer’s Disease. Here, we report that loss of *sfpq* leads to premature termination of multiple transcripts due to widespread activation of previously unannotated cryptic last exons (CLEs). These CLEs appear preferentially in long introns of genes with neuronal functions and dampen gene expression outputs and/or give rise to short peptides interfering with the normal gene functions. We show that one such peptide encoded by the CLE-containing *epha4b* mRNA isoform is responsible for neurodevelopmental defects in the *sfpq* mutant. The uncovered CLE-repressive activity of SFPQ is conserved in mouse and human, and SFPQ-inhibited CLEs are found across ALS iPSC-derived neurons. These results greatly expand our understanding of SFPQ function and uncover a new gene regulation mechanism with wide relevance to human pathologies.

## Introduction

Neurons are highly polarized cells with specialized compartments that must be able to respond to growth cues as well as to form and modify their synapses in an activity-dependent manner. Each compartment of a neuron is able to achieve functional specificity by maintaining a unique proteome (Holt & Schuman, 2013; Hanus & Schuman, 2013; Cagnetta *et al*, 2018). Protein localization in neurons has been shown to be driven largely by RNA transportation and local translation (Zappulo *et al*, 2017), suggesting that neuronally-expressed genes must have special regulatory mechanisms to ensure proper transcription, localization, and translation of each RNA. Indeed, RNAs from neuronal tissue are regulated by a complex array of alternative splicing, intron retention, and alternative cleavage and polyadenylation (Mauger *et al*, 2016; Traunmüller *et al*, 2016; Furlanis *et al*, 2019; Iijima *et al*, 2019; Taliaferro *et al*, 2016; Ciolli Mattioli *et al*, 2019; Guvenek & Tian, 2018; Tushev *et al*, 2018).

Splicing Factor Proline/Glutamine Rich (SFPQ) is a ubiquitously expressed RNA binding protein of the DBHS family with diverse roles in alternative splicing, transcriptional regulation, microRNA targeting, paraspeckle formation, and RNA transport into axons (Patton *et al*, 1993; Dye & Patton, 2001; Kim *et al*, 2011; Cosker *et al*, 2016; Bottini *et al*, 2017; Mora Gallardo *et al*, 2019; Takeuchi *et al*, 2018; Knott *et al*, 2016). Inactivation of the *sfpq* gene causes early embryonic lethality in mouse and zebrafish as well as impaired cerebral cortex development, reduced brain boundary formation, and axon outgrowth defects (Lowery *et al*, 2007; Thomas-Jinu *et al*, 2017; Takeuchi *et al*, 2018; Saud *et al*, 2017). In humans, *sfpq* mutations have been linked to neurodegenerative diseases such as Alzheimer’s, ALS, and FTD, and SFPQ interacts with the ALS-associated RNA binding proteins TDP-43 and FUS (Ke *et al*, 2012; Wang *et al*, 2015; Ishigaki *et al*, 2017; Luisier *et al*, 2018; Tyzack *et al*, 2019; Lu *et al*, 2018).

While SFPQ is known to play a role in alternative splicing, only a few RNA targets of SFPQ have been identified. Intriguingly, SFPQ has opposing effects on splicing, depending on the target: it represses inclusion of exon 10 of tau and exon 4 of CD45, but conversely it promotes inclusion of the N30 exon of non-muscle myosin heavy-chain II-B (Ray *et al*, 2011; Ishigaki *et al*, 2017; Heyd & Lynch, 2010; Yarosh *et al*, 2015; Kim *et al*, 2011). In addition to its role in splicing, SFPQ has been shown to be part of the 3’-end processing complex, where it enhances cleavage and polyadenylation at suboptimal polyadenylation sites (Hall-Pogar *et al*, 2007; Rosonina *et al*, 2005; Shi *et al*, 2009). The mechanisms by which SFPQ regulates mRNA processing are still unclear, however, and more work is necessary to understand its contribution to normal and pathological cell states.

To understand the molecular functions of SFPQ in developing neurons, we performed an RNA-seq analysis of *sfpq* homozygous null mutant zebrafish embryos at 24 hpf, the stage of phenotypic onset. Our results reveal a novel role for the protein: loss of SFPQ causes premature termination of transcription as a result of previously unannotated pre-mRNA processing events that we refer to as Cryptic Last Exons (CLEs). Here we describe the formation of CLEs and show that not only do the truncated transcripts act as a form of negative regulation of gene expression levels, but they also directly contribute to the *sfpq* pathology. This function of SFPQ is conserved across vertebrates and may be implicated in human SFPQ-mediated disease states.

## Results

### Identification of the SFPQ-dependent splicing regulation program

To examine the effect of SFPQ on gene expression and RNA splicing, we analyzed total RNA extracted from 24 hpf *sfpq*^-/-^ zebrafish embryos and their heterozygous or wildtype siblings by RNA sequencing (RNA-seq). Differential gene expression analysis using Cufflinks RNA-seq workflow (Trapnell *et al*, 2012) uncovered 189 genes that were upregulated and 1044 genes that were downregulated in the mutant samples by a factor of at least 1.3-fold with q≤0.05 (Figure 1a). These results are consistent with our previous microarray study, which showed the vast majority of genes with differential expression in *sfpq*^-/-^ embryos as being downregulated (Thomas-Jinu *et al*, 2017). Gene ontology (GO) analysis of the new dataset, using total transcribed genes as a background gene set, showed enrichment for neuron-specific terms, including neuronal differentiation and axon guidance (Tables S1 and S2). Using Cufflinks’ differential isoform switch analysis, we identified 112 genes with significant change in the relative expressions of splice variants in the mutants (q≤0.05; Table S3). GO analysis of these regulated transcripts again showed an over-representation of neuron-specific terms including axonogenesis, axon guidance, and dendrite formation (Table S4). Surprisingly, thorough comparison and annotation of these transcripts revealed that 46% of these genes express a splice variant containing a cryptic alternate last exon, not annotated in the zebrafish assembly (Figure S1a). To verify this, we analyzed the dataset with Whippet (Sterne-Weiler *et al*, 2018), a tool that sensitively detects changes in the usage of alternative exons and additionally allows quantitation of gene expression changes. Whippet also uncovered a high proportion of downregulated genes in *sfpq*^-/-^ embryos (Figure 1b). More importantly, the analysis confirmed that splicing of alternate last exons is the most abundant (18.5%) category of SFPQ-regulated splicing events (Figure 1c). Systematic classification of these exons into “known” and “cryptic” events corroborates that the majority (113 out of 157) of these last exons have not been previously annotated (Figure 1D, Table S5). These last exons were expressed from 106 genes, 25 of which were also detected by the Cufflinks pipeline (Figure S1b). We refer to this pervasive splicing defect, in which the transcript undergoes premature termination after the inclusion of a cryptic exon, as Cryptic Last Exons (CLEs) (Figure 1e).

**Figure 1:**
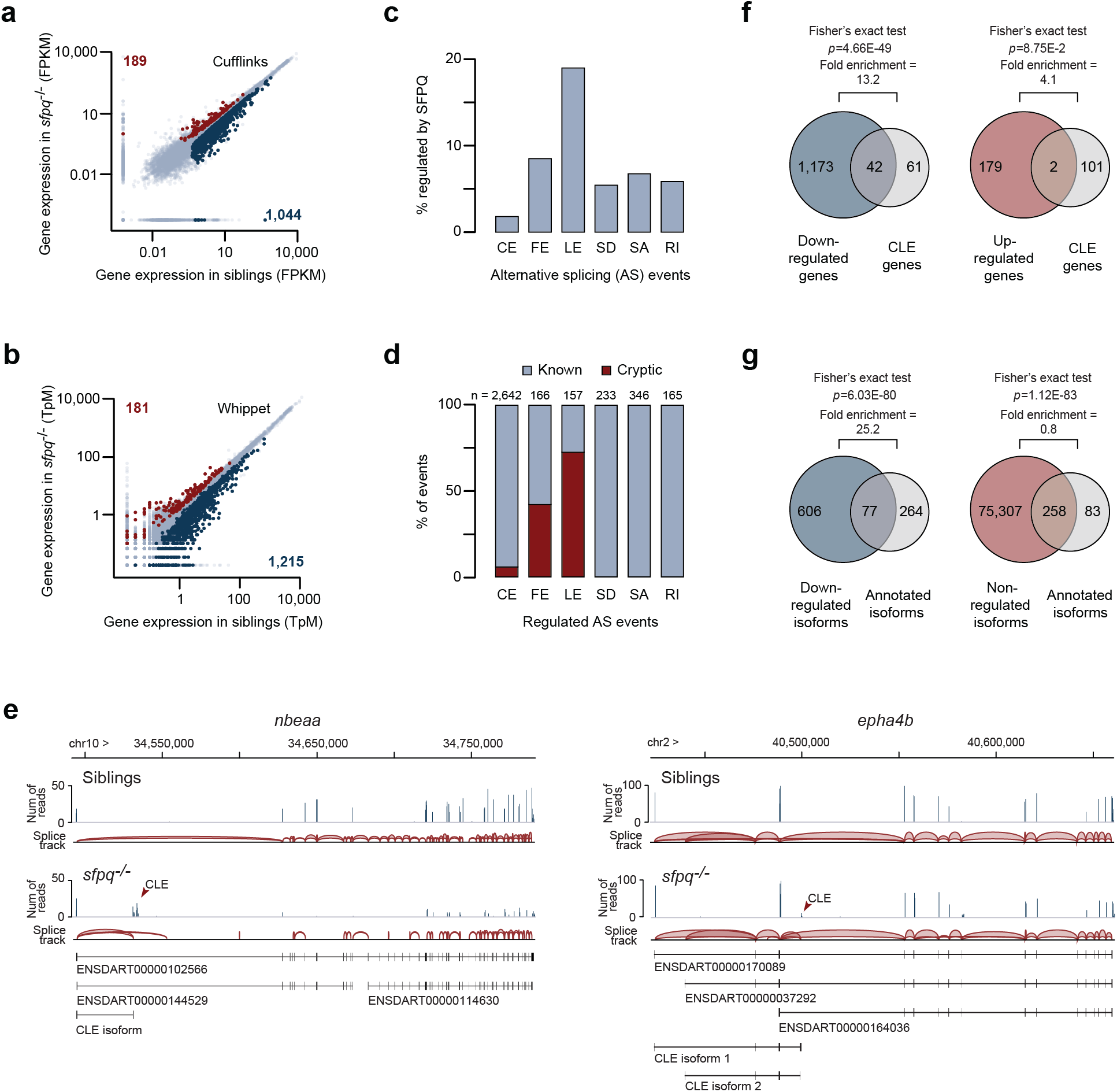
SFPQ regulates the formation of cryptic last exons (CLEs) **a-b,** Scatter plot showing expression values of genes in sfpq−/− and siblings, analyzed using Cuffiinks (a) or Whippet (b) pipelines. **c,** Alternative last exon splicing is highly regulated by SFPQ. CE: cassette exon, FE: first exon, LE: last exon, SD: splice donor, SA: splice acceptor, RI: retained intron. **d,** Majority of SFPQ-regulated last exon events are cryptic. **e,** Sashimi plots showing example CLE formation in nbeaa and epha4b. Top tracks: plot of read coverage from siblings (upper) and sfpq−/− (lower). Bottom tracks: isoforms discovered for each gene. **f,** Genes expressing CLE-containing isoforms tend to be down-regulated in sfpq−/−. **g,** Normal long isoforms (annotated isoforms) from CLE-expressing genes tend to be down-regulated in sfpq−/−.

### The use of CLEs inversely correlates with expression of full-length transcripts

Of the 106 CLE-expressing genes, 97% exhibited increased splicing of CLEs in *sfpq*^-/-^ (Table S5). Notably, more than half of these genes were downregulated in mutants (~13 fold enrichment over the number of genes expected by chance; Fisher’s exact test p=4.66 × 10^−49^) indicating concurrent alterations in expression level and splicing for these genes (Figure 1f). In line with this finding, the full-length (non-CLE) isoforms from these genes showed an even stronger enrichment for the downregulation effect (exceeding the expectation ~25-fold; Fisher’s exact test p=6.03 × 10^−80^) (Figure 1g). To verify these results, we performed RT-qPCR on five selected CLE-containing genes: *nbeaa*, *gdf11*, *epha4b*, *trip4*, and *b4galt2*. In all cases, cryptic exons showed a substantial increase in expression level in *sfpq*^-/-^ mutants compared to siblings (Figure S1c-g). Additionally, we detected a strong downregulation of the full-length isoforms in four of the five genes, suggesting that the loss of *sfpq* causes upregulation of the CLE isoforms at the expense of their normal counterparts (Figure S1d-g). These results argue that SFPQ is required to repress CLE splicing in order to maintain stable gene expression.

### CLEs tend to occur in long introns and show evidence of interspecies conservation

In order to understand under what conditions CLEs form, we examined CLE-containing introns and compared them to all other introns from the same genes. We first asked where CLE-containing introns are found within their genes and found no bias (Figure 2a). However, when we ranked the introns by length, we found that CLEs are frequently located in the longest intron of the gene (Figure 2b). CLE-containing introns are also significantly longer than the average intron size in the entire zebrafish transcriptome (Figure 2c). Consistent with these results, CLE-containing genes are significantly longer than average zebrafish genes (Figure S2). Within the intron, location of the CLE is biased toward the 5’ end, with most appearing approximately 22.4% (95% confidence interval of 18.1% to 26.7%) of the way into the intron (Figure 2d). The distance between the CLE and the upstream exon is generally <10 kb (Figure 2e). We next asked whether the sequences within and neighboring these CLEs are conserved. To this end, we calculated the mean conservation scores of 1 kb sequences (sliding window, 1 bp steps) along these CLE-containing introns, using the PhastCons analysis method (Siepel *et al*, 2005). Our analyses showed that sequences containing CLEs tend to have higher conservation scores as compared to sequences within the same intron that do not contain CLEs (Figure S2b). In fact, 18% of these sequences displayed a mean PhastCons score of at least 0.5 (as opposed to 12% of non-CLE sequences; Fisher’s exact test: 5.71 × 10^−51^) (Figure S2c). Next, we calculated the mean base conservation scores of each CLE together with 250 bp flanking sequences (Figure 2f and S2d). Although only 35% of the CLEs had a PhastCons score of at least 0.5, the sequences near its 3’ acceptor site showed the highest conservation (Figure 2f). Together, these data indicate that CLEs are often found close to the 5’ ends of very long introns and that at least some of these exons are evolutionarily conserved.

**Figure 2:**
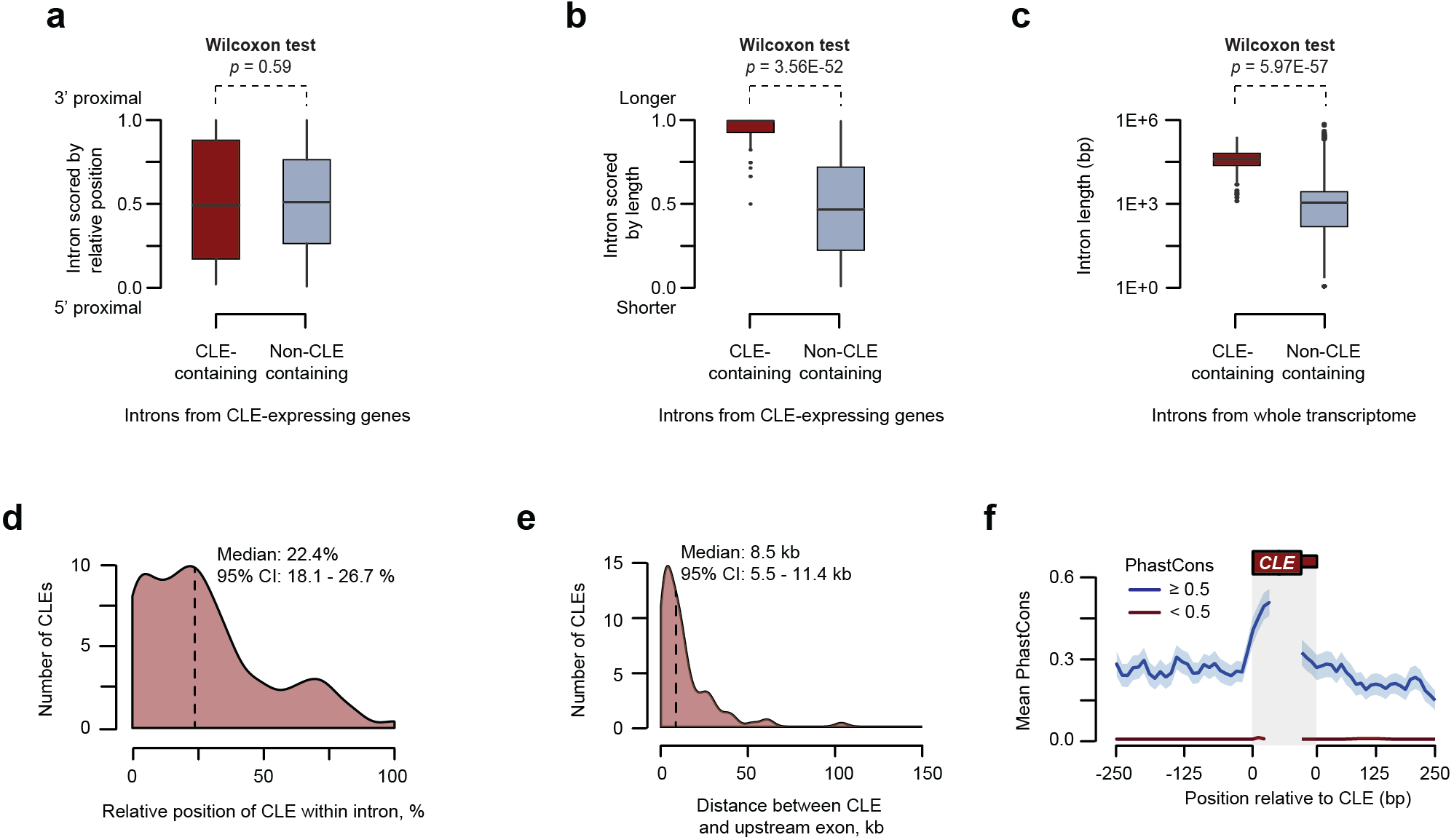
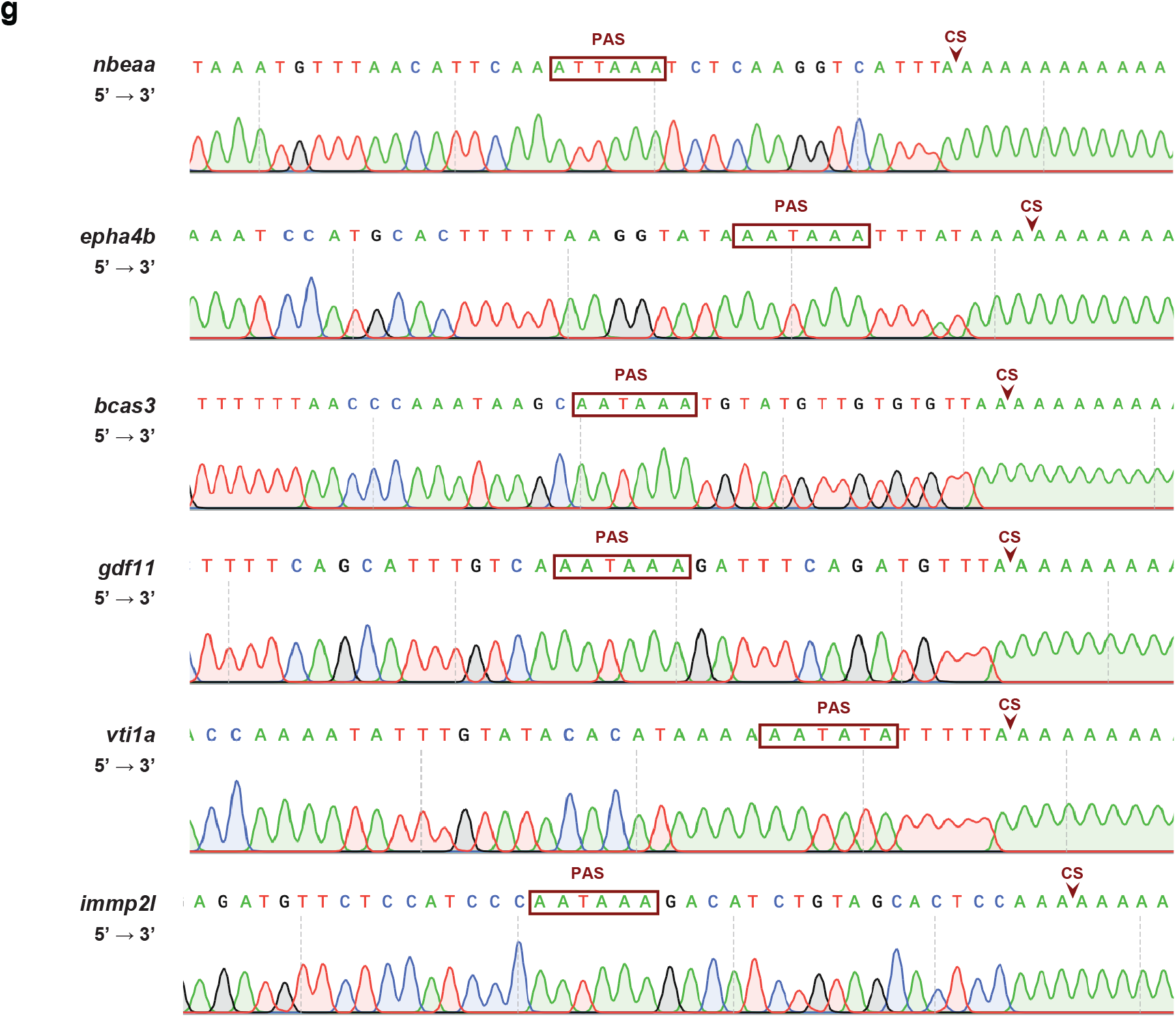
Molecular properties of CLEs. **a-b,** Introns from CLE-expressing genes were scored by its relative position (a) and by its relative length (b), and the distribution of these scores were plotted. Note that introns containing CLE tend to be long and sparsely distributed. **c,** CLE-containing introns are longer than average intrans. Length of CLE-containing introns is compared to all other introns from the zebrafish transcriptome. **d,** CLEs tend to be found closer to the 5’ end of its intron. **e,** CLEs are found within 10 kb of the upstream constituitive exon. **f,** Line-plot showing the conservation score of sequences surrounding conserved (blue) and non-conserved (red) CLEs. 280 bp of surrounding intron/CLE junction sequence (250 bp intron and 30bp exon) were binned into 10 bp windows and the mean PhastCons score for each bins were shown (± SEM). **g,** Sanger sequencing of 3’RACE PCR products of CLE isoforms. PAS hexamers are shown within red boxes and the predicted cleavage site are marked by arrowheads.

### CLE-terminated transcripts are cleaved and polyadenylated at the 3’ end

Our data thus far suggests that these cryptic transcripts are stably expressed and detectable using RNA-seq and RT-qPCR techniques. Sequence analyses revealed that 63% of these transcripts contain an open reading frame predicted to express truncated peptides with missing C-terminal domains (Table 6). To test if CLE transcripts are polyadenylated, we performed 3’ RACE on six CLE-containing transcripts: *bcas3*, *epha4b*, *gdf11*, *immp2l*, *nbeaa*, and *vti1a*. We found that all six showed elements of strong polyadenylation sites (Shi & Manley, 2015): four of the six exons had canonical AAUAAA hexamers just upstream of the cleavage site, while the other two had common one-base substitutions of AUUAAA and AAUAUA. In addition, five of the six contained downstream GUGU sequences, while two also had an upstream UGUA. Although none of the exons had a canonical CA sequence directly 5’ of the cleavage site, overall the cryptic exons displayed strong polyadenylation sequences.

### SFPQ directly binds to sequences adjacent to CLEs

The accumulation of CLE-terminated transcripts in *sfpq* mutants raises the question of whether SFPQ represses CLEs in a direct manner. SFPQ binds promiscuously to a wide range of RNA sequences (Yarosh *et al*, 2015; Knott *et al*, 2016), making binding prediction difficult. Using a binding motif produced by a recent *in vitro* study (Ray *et al*, 2013), however, we found a significant enrichment in predicted SFPQ binding sites upstream of cryptic exon sequences compared to control last exons (Figure 3a). To validate this, we purified SFPQ-RNA complexes in 24 hpf embryos using standard CLIP protocol and quantified the relative amount of bound CLE RNA fragments using RT-qPCR. Our results confirmed that SFPQ binds either within the CLE or in adjacent 5’ or 3’ intronic regions of at least three CLE transcripts (Figure 3b-d). These results support the idea that SFPQ directly binds to region surrounding CLEs to regulate their inclusion.

**Figure 3:**
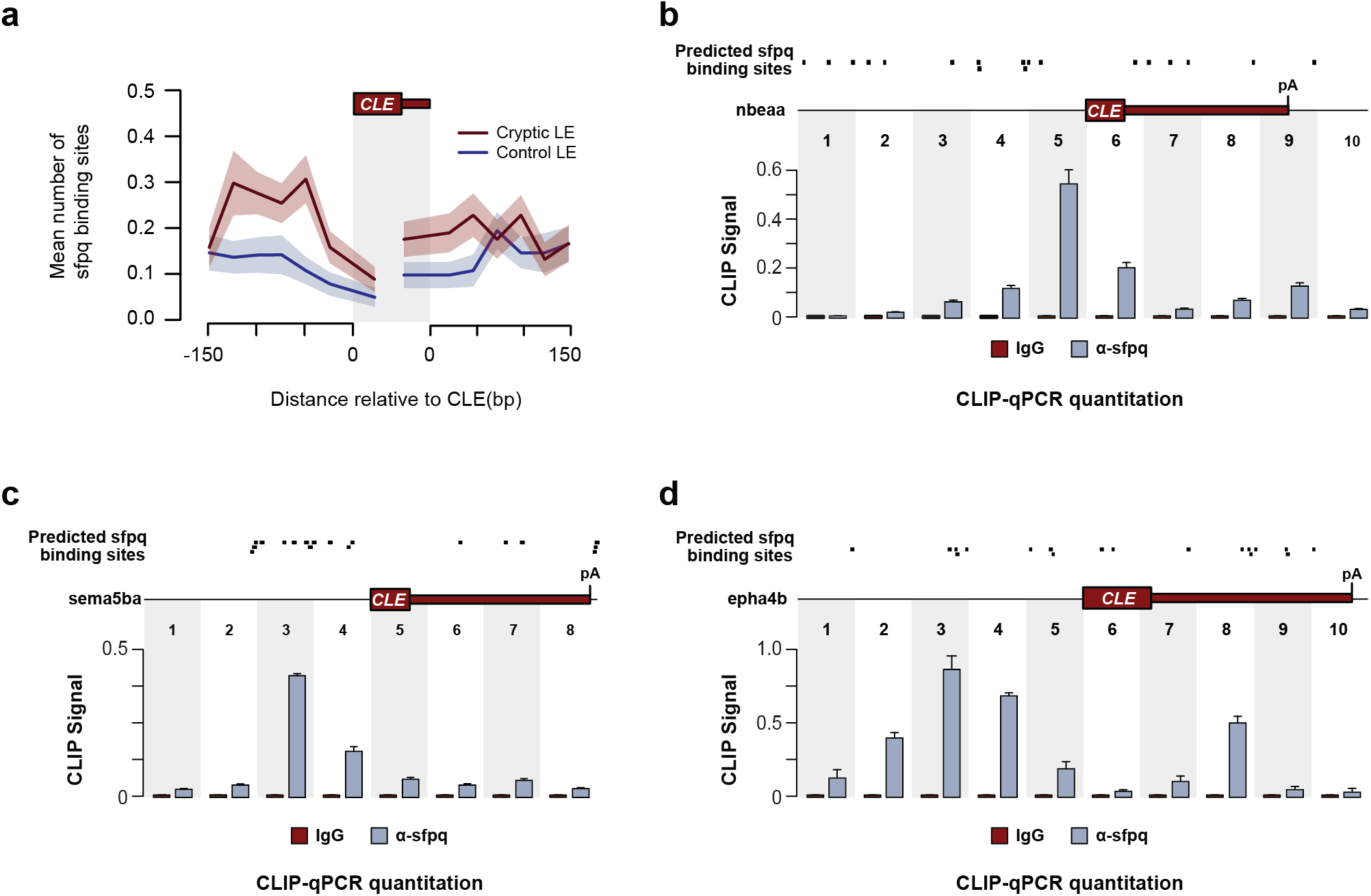
SFPQ directly binds to RNA adjacent to CLE sequences. **a,** Line plot showing the distribution of predicted SFPQ-binding sites surrounding CLEs (red) and constitutive last exons of each CLE-containing gene(blue). 200 bp of surrounding intron/CLE junction sequence(150 bp intron and 50bp exon) were binned into 50 bp windows and the mean number of predicted motifs were shown(± SEM). **b-d,** Top: Location of SFPQ binding motifs predicted using MEME suite. Bottom: RT-qPCR quantitation showing the relative enrichment of SFPQ-interacting regions surrounding CLEs. Abundance of SFPQ- or lgG(control)-crosslinked RNAs were normalized to input and the mean value from three replicates were shown(± SD).

### CLE<u>s</u> can dampen the expression of full-length transcripts

The reciprocal relationship between CLEs and the abundance of full-length transcripts (Figure 1f) suggests that these exons may act as negative regulators of gene expression. If production of CLE transcripts is a mechanism for down-regulating the normal full-length transcripts, then eliminating the cryptic exon in *sfpq*^-/-^ mutants should rescue their expression. To test this possibility, we used the gene *b4galt2* as case study, as it shows a very strong loss of expression of its three normal isoforms in the mutant (Figure S1f). We used CRISPR/Cas9 to delete the *b4galt2* CLE, injecting Cas9 along with two guide RNAs that targeted directly upstream of the cryptic exon and at the 3’ end of the exon (Figure 4a). Injected founder embryos (crispants) will show mosaicism, so a complete loss of the cryptic exon would not be expected in every cell of the embryo. Despite mosaicism, PCR analysis of the “crispants” showed a strong deletion band for six out of eight tested embryos (Figure 4a). Encouraged by the high efficiency of the gRNAs, we performed RT-qPCR on pooled injected *sfpq*^-/-^ embryos to measure the expression levels of the normal *b4galt2* transcripts and saw a significant rescue of the longer transcripts compared to the uninjected *sfpq*^-/-^ control (Figure 4a). This result did not hold true with two other CLEs we deleted (example *gdf11* CLE, Figure S4a). We concluded that CLEs can regulate expression levels of at least some of the genes containing these exons.

**Figure 4:**
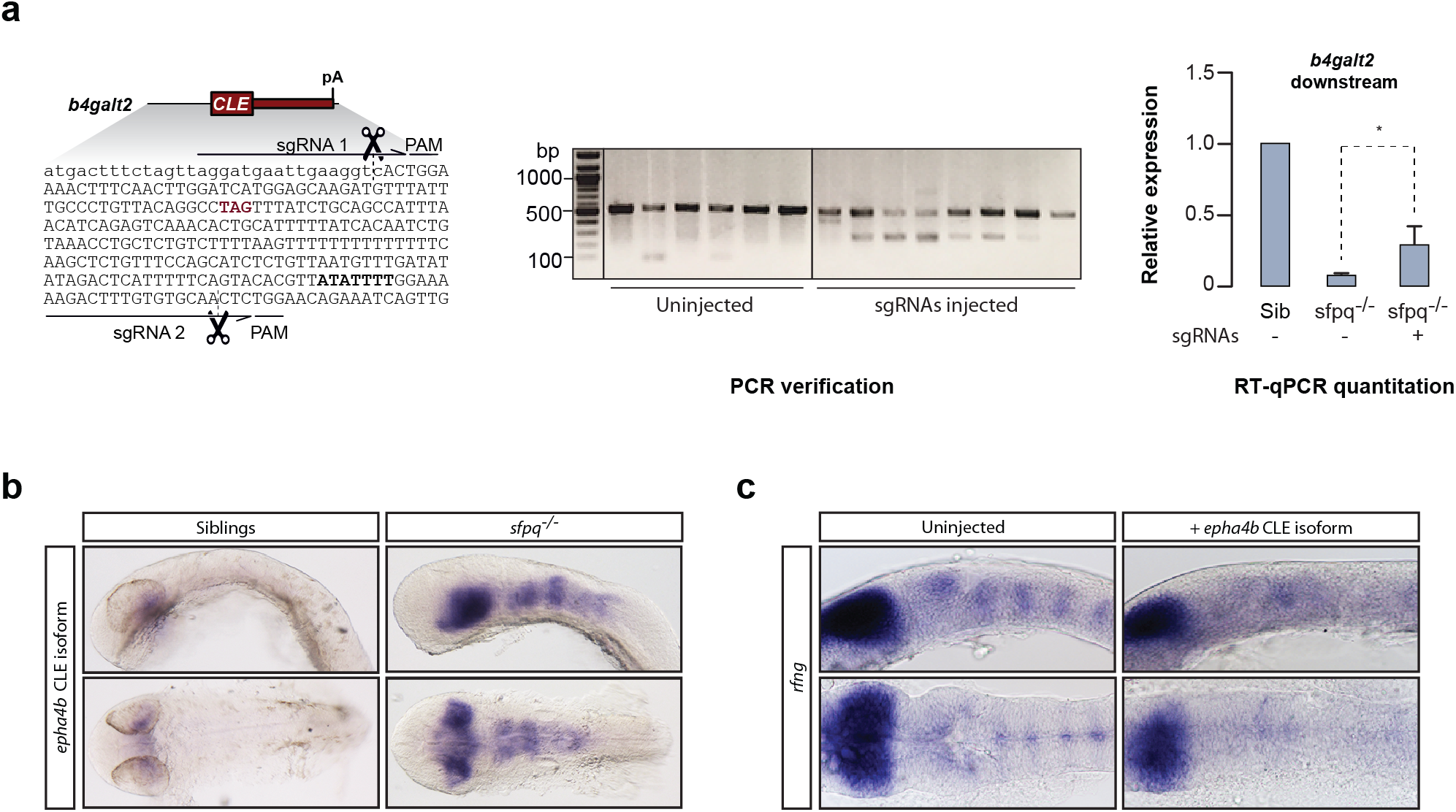
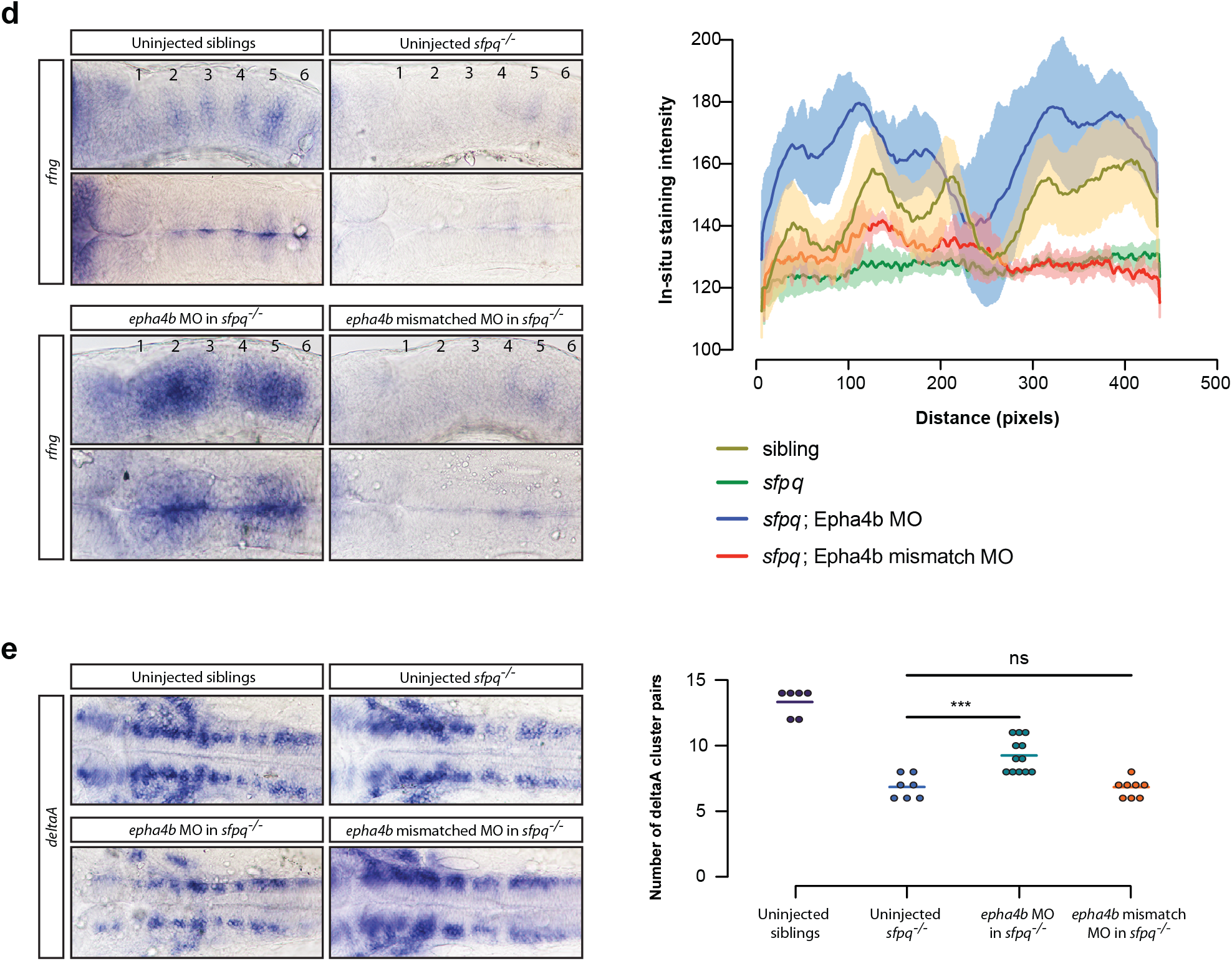
CLE formation is functionally relevant. **a,** Deletion of the b4galt2 CLE using CRISPR/Cas9 rescues expression of downstream exons. Left: cut sites of the b4galt2 sgRNAs. CLE is indicated by capital letters. Center: PCR verification of Cas9 cleavage after injection of sgRNAs. Right: RT-qPCR quantitation of the relative expression of the downstream b4galt2 exons in sfpq−/− embryos compared to siblings. **b,** in-situ hybridization of the epha4b CLE at 24 hpf, displaying strong expression in the midbrain and hindbrain of sfpq−/− embryos. **c,** in-situ hybridization of rfng shows rhombomere boundary defects at 22ss after injection of the epha4b cryptic transcript into WT embryos **b-d,** Upper: lateral view. Lower: dorsal view. **d,** Left: in-situ hybridization of rfng shows rhombomere boundary defects of sfpq−/− embryos are rescued by injection of the epha4b cryptic splice junction morpholino but not a mismatch morpholino. Rhombomere boundaries are numbered. Right: quantification of staining in rhombomeres in three lateral view samples for each condition **e,** Left: in-situ hybridization of DeltaA shows a loss of discrete neuronal clusters in sfpq−/− which is rescued by injection of the epha4b cryptic splice junction morpholino but not a mismatch morpholino. Right: Quantification of number of DeltaA clusters in each condition. **b-d,** Upper: lateral view. Lower: dorsal view.

### Truncated protein derived from CLE-containing *epha4b* transcripts accounts for the boundary defects in *sfpq*^-/-^ brain

In addition to affecting the expression levels of normal isoforms, CLE transcripts could impact the *sfpq* phenotype through aberrant functions of the truncated RNAs or the short peptides they produce. We focused on the candidate gene *epha4b*, which expresses a CLE-containing short mRNA in *sfpq* null embryos, while showing no change in expression of the normal transcripts (Figure S1c). This gene is one of two zebrafish paralogues of the human ALS-associated gene Epha4 (Van Hoecke *et al*, 2012; Wu *et al*, 2017), coding for a protein-tyrosine kinase of the Ephrin receptor family known to regulate hindbrain boundary formation (Cooke *et al*, 2005; Kemp *et al*, 2009). Truncated forms of EPH receptors have been shown to act as dominant negatives by competing with full-length versions of the protein for ligand binding (Smith *et al*, 2004). The predicted peptide produced by the *epha4b* CLE-containing short transcript would contain the ligand binding domain but not the transmembrane and intracellular domains and thus would be predicted to be a dominant negative (Table S6).

To assess possible effects of the shortened *epha4b*, we first performed an *in-situ* hybridization using a probe for the cryptic exon. We found that in *sfpq*^-/-^ embryos, but not in siblings, the *epha4b* CLE was expressed strongly in the midbrain and hindbrain (Figure 4b), where the gene is normally transcribed at that developmental stage. We then tested whether, in wildtype fish, injection of the CLE transcript would induce defects in the midbrain or hindbrain. Using the early hindbrain boundary marker *rfng*, we found that injection of the short *epha4b* transcript did not affect formation of the midbrain but did cause a loss of hindbrain rhombomere boundaries similar to that seen in the *sfpq*^-/-^ mutant (Figure 4c).

We then asked whether repressing the *epha4b* CLE in *sfpq*^-/-^ embryos could rescue the *sfpq* hindbrain defect. We used a splice junction morpholino (MO) that targeted the 3’ splice acceptor site of the CLE to prevent the cryptic exon from being used in *sfpq* mutants. Although MOs frequently have off-target effects, those effects are generally the opposite of what we would expect to see from a rescue (i.e. increased cell death and off-target phenotypes, never rescue of phenotypes). However, as MOs have been shown to have some phenotypic effects on the hindbrain (Gerety & Wilkinson, 2011), we used mismatch controls to ensure that our results were specific to the *epha4b* CLE splice-MO. We tested the MO efficiency using RT-PCR with primers both within the cryptic exon and across the exon junction (Figure S4b). We then examined the effects of the MO on hindbrain development using both the boundary-specific *rfng* marker (Figure 4d) and the pan-neuronal marker DeltaA (Figure 4e). We saw that the CLE splice junction MO, but not the mismatch control, rescued formation of rhombomere boundaries in *sfpq*^-/-^ mutants. Taken together, these results indicate that the hindbrain boundary defect in *sfpq*^-/-^ embryos can be explained by the dominant-negative effects of the *epha4b* CLE transcript.

### Repression of CLEs by SFPQ is conserved across vertebrates and relevant to human neuropathologies

As our analysis of the *sfpq* loss-of-function phenotype was performed solely in zebrafish, we wondered whether SFPQ repressed CLEs in other organisms. Accordingly, we turned to publicly available RNA-seq datasets from *sfpq* loss-of-function experiments. A conditional mouse knock-out model (Takeuchi *et al*, 2018) inactivated *Sfpq* in the cerebral cortex. Examining mouse orthologs of zebrafish CLE-containing genes, we were able to identify CLE formation in mouse *Sfpq*-null brains for *Epha4b*, *Cpped1*, *Fam172a*, and *Exoc4* (Figure 5a and S5). Overall, we identified 144 instances of upregulation of CLEs in the cortical *Sfpq* knockout (Table S7). Examination of the CLE-containing introns showed results similar to those for the zebrafish CLEs: the CLE-containing introns have a bias towards appearing earlier in the gene, they are often embedded within the largest intron in a gene, and their host introns are significantly larger than the average mouse intron (Figure 5b).

**Figure 5:**
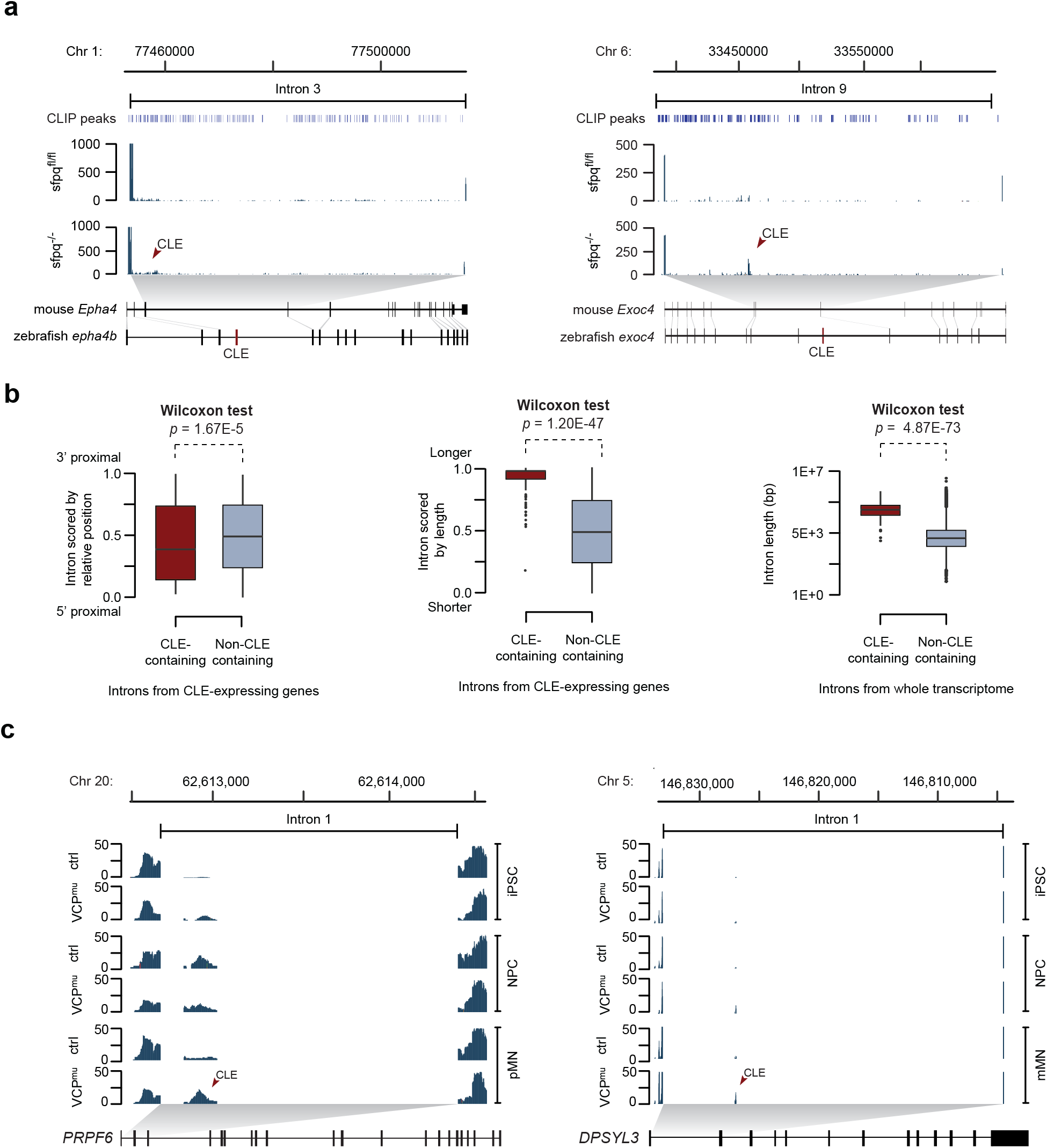
The CLE-repressing function of SFPQ is conserved in mouse and human. **a,** Meta-analysis of RNA-seq and CLIP-seq dataset from conditional Sfpq knockout mice for cryptic ast exons. Top: distribution of Sfpq CLIP peaks within the CLE-containing intron. Middle: tracks 5howing read coverage plots and “sashimi” plots from Sfpqfl/fl and Sfpq−/− mice. Bottom: exon architecture of orthologous CLE-expressing genes. Homologous regions between orthologues are 5hown as connecting lines. **b,** lntrons from mouse CLE-expressing genes were scored by its relative position (left) and by its relative length (mid), and the distribution of these scores were plotted. Note that introns containng CLE tend to be long and sparsely distributed. Right: CLE-containing introns are longer than 3verage introns. Length of CLE-containing introns is compared to all other introns from the mouse transcriptome. **c,** Representative RNA-seq coverage plots from ALS-derived iPSC dataset of CLEs up-regulated in VCPmu samples.

As SFPQ has been recently linked with ALS in human, we also examined RNA-seq results from iPSCs derived from ALS patients, which show loss of nuclear SFPQ expression (Luisier *et al*, 2018). In total, we found 76 CLE events up-regulated in ALS-mutant backgrounds across the neuronal differentiation stages (Table S8). This is probably an underestimation since the sequencing depth in this dataset was somewhat lower than that in the mouse knockout study. Interestingly, CLEs spliced from PRPF6 and DPYSL3 genes showed consistent up-regulation in three time-points (Figure 5c). The latter gene is involved in positive regulation of axon guidance and genetic variants of this gene have been previously implicated in ALS patients (Blasco *et al*, 2013). These results indicate that CLE repression is a conserved function of SFPQ, and that CLE-dependent short transcripts may have a substantial impact on SFPQ-mediated disease states.

## Discussion

Our study uncovers a critical role of SFPQ in repression of cryptic last exons (CLEs). We show that truncated transcripts appearing as a result of increased use of CLEs are functionally relevant both as regulators of gene expression output and as a source of interfering protein isoforms. Moreover, the CLE-repressing function of SFPQ is conserved in mouse and human, indicating an important developmental role, with implications for human pathology.

### Mechanism of CLE formation

The presence of strong polyadenylation sites in CLE sequences suggests that the paucity of CLE-containing isoforms under normal physiological conditions is due to active suppression of CLE cleavage/polyadenylation or/and splicing. Our CLIP-seq experiments provide evidence for SFPQ binding within or directly adjacent to CLEs. Moreover, the bias of CLEs towards the 5’ end of long introns is consistent with previous analyses of SFPQ localization on RNA (Takeuchi *et al*, 2018). These data argue that SFPQ may play a direct role in repressing cryptic exon formation. However, further work will be required to distinguish between suppression of splicing versus blocking of the polyadenylation site.

The relationship between SFPQ and CLEs extends our understanding of the regulation possibilities afforded by long introns. Indeed, long introns have been previously shown to control gene expression through interplay between premature cleavage/polyadenylation and the U1 snRNP-dependent antitermination mechanism known as “telescripting” (Langemeier *et al*, 2013; Oh *et al*, 2017; Venters *et al*, 2019; Kainov & Makeyev, 2020). SFPQ-mediated CLE repression also operates in long introns (Figure 2b) but, unlike telescripting, CLE involves definition of a last exon possibly via interactions between the U2 snRNP and U2AF with the cleavage/polyadenylation machinery (Martinson, 2011). Moreover, inactivation of U1 often promotes cleavage/polyadenylation relatively close to the 5’ end of the gene, whereas CLEs do not show such a gene location bias.

Long introns have been also shown to be subject to recursive splicing (RS), a multistep process promoting accuracy and efficiency of intron excision (Sibley *et al*, 2015; Blazquez *et al*, 2018). Like CLEs, RS-sites appear primarily in long introns in genes with neuronal function. RS-sites initially produce an RS-exon that is spliced to the upstream exon prior to being excised at the subsequent round of splicing reactions. However, RS-exons do not contain polyadenylation sequences, so inclusion of the exon would not lead to truncation of the transcript. In addition, recursive splicing creates a stereotypical saw-tooth pattern of RNA-seq reads, which is not seen in the *sfpq*^-/-^ RNA-seq data set. Therefore, SFPQ and CLEs provide a distinct regulation modality compared to telescripting and recursive splicing.

### Pathology of cryptic transcripts

A notable feature of the SFPQ-repressed CLEs is the detrimental effect that they have on the function of their host genes. We previously showed that loss of *sfpq* leads to an array of morphological and neurodevelopmental abnormalities in zebrafish embryos, including loss of brain boundaries and altered motor axon morphology (Thomas-Jinu *et al*, 2017). However, the mechanism by which those abnormalities formed was unresolved. Here, we found that CLEs contribute to at least one aspect of the *sfpq* phenotype: the dominant negative *epha4b* truncated transcript induces hindbrain boundary defects. Moreover, a subset of the identified CLE-dependent short transcripts identified in *sfpq*^-/-^ is predicted to affect axon growth and connectivity.

While CLE formation is clearly detectable under pathological conditions of loss of *sfpq*, our data do not preclude the possibility of CLEs being expressed under non-pathological conditions. Although the CLEs are not annotated in the current zebrafish, mouse, and human genomes, it is possible that they may be regulated in a spatio-temporal manner such that they only appear in specific tissues and/or at specific developmental time points. Indeed, this possibility is supported by the relatively low expression of SFPQ in non-neuronal tissue (Thomas-Jinu *et al*, 2017; Lowery *et al*, 2007), and by low-level detection of the *epha4b* CLE transcript in siblings by PCR (Figure S4B). Early termination of long pre-mRNAs has been shown to be a developmentally controlled regulatory mechanism: the RNA-binding protein Sex-lethal promotes the formation of truncated transcripts during short nuclear cycles in *Drosophila* (Sandler *et al*, 2018), and downregulation of the cleavage and polyadenylation factor PCF11 during differentiation of mouse C2C12 myoblast cells suppresses intronic polyadenylation to promote long gene expression (Wang *et al*, 2019). Further examination of CLE expression in wildtype animals across development may identify possible role of these truncated transcripts in normal tissues.

### Cryptic exons in neurodegenerative disease

Neurodegenerative diseases such as Alzheimer’s, ALS, and FTD are frequently characterized by altered localization and function of splicing factors (Tyzack *et al*, 2019; Ling *et al*, 2013; Neumann *et al*, 2006; Nag *et al*, 2018). The ALS-associated proteins transactivation response element DNA-binding protein 43 (TDP-43) and fused in sarcoma (FUS) regulate alternative splicing and alternative polyadenylation (Ishigaki *et al*, 2012; Masuda *et al*, 2016; Deshaies *et al*, 2018; Melamed *et al*, 2019; Klim *et al*, 2019; Ling *et al*, 2015). TDP-43 has been shown to act as a repressor of cryptic exons, a minority of which contain polyadenylation sites and thus would form CLEs (Ling *et al*, 2015). Stathmin-2 is one of the latter and rescue of its normal full-length expression in TDP-43-knockdown cell culture improves axonal growth in this model (Melamed *et al*, 2019; Klim *et al*, 2019), indicating that CLEs are pathogenic across various splicing protein-dependent pathologies. These findings place CLEs at the center of priority for understanding molecular mechanisms of neurodegenerative diseases and developing new ways to diagnose and treat these increasingly prevalent disorders.

## Acknowledgements

This work was supported by the Biotechnology and Biological Sciences Research Council (BB/P001599/1 to CH BB/R001049/1 to E.V.M.); the European Commission (Project ID 734791; E.V.M); the Wellcome Trust (Tech Dev. WT093389 and Equipment WT094819 to CH) and EMBO (fellowship ALTF 1530-2015 to PMG). We thank Richard Poole for help with the TopHat pipeline.

## Materials and Methods

### Zebrafish husbandry

Zebrafish (*Danio rerio*) were reared in accordance with the Animals (Scientific Procedures) Act 1986. Fish were maintained on a 14 hr light/10 hr dark cycle at 28**°**C. Embryos were cultured in fish water containing 0.01% methylene blue to prevent fungal growth. Wildtype fish were AB strain from the Zebrafish International Resource Center (ZIRC), while *sfpq* null mutants were *sfpq*^*kg41*^ (Thomas-Jinu *et al*, 2017).

### RNA-seq

RNA was extracted from 24 hpf *sfpq*^-/-^ embryos and their heterozygous or homozygous wildtype siblings using the RNeasy Mini Kit (Qiagen). RNA was sequenced using the Illumina HiSeq 2500 with 50bp paired-end reads.

### 3’ RACE

RNA was extracted from 24 hpf *sfpq*^-/-^ embryos using the RNease Mini Kit (Qiagen). Reverse transcription was performed using the 3’ RACE System for Rapid Amplification of cDNA Ends (ThermoFisher). cDNA was amplified in two subsequent PCR reactions, using the adapter primer as a reverse primer and the following primers as forward primers:

*nbeaa:* AGAGAGGGACCGTGTAGAC, AAGGCACATCGAGCCCATATTG
*epha4b:* ATGGCAACCCTTTGGATTTATCCG, CTACTGTCAGGCTGTTTCGG
*bcas3:* CGCTGCATGTCAGCTTCAC, CTTCAGGAAACTGACAAACGCGAG
*gdf11:* GGGAGCATTATAGGCATCGGTAC, TGGCTTCAGAGCGAGTCATAG
*vti1a:* CGTCAATAAGAAGCAGACACAAGCAAC, GATTTTGTGGTCACATTTTCGTG
*immp2l:* CGACGCACAGCACGTACATAAG, GCGACATTGTGTCAGTTTTAACATC

### CLIP-qPCR

Dechorionated 24 hpf wildtype fish were irradiated (twice at 0.8 J/cm^2^, 254nm) and deyolked using high calcium Ringer’s solution (116 mM NaCl, 2.9 mM KCl, 10 mM CaCl2, 5 mM HEPES, pH 7.2) with 0.3 mM PMSF and 1 mM EDTA. After several washes, embryos were lysed using PXL buffer (0.1% SDS, 0.5% deoxycholate, 0.5% NP-40) and homogenized using a plastic pestle. Lysates were treated with 10 μL diluted RNAseI (1:500 dilution; Thermo Fisher) and 2 μL Turbo DNase (Thermo Fisher) at 37°C for 3 minutes on a shaking incubator. Protein-RNA complexes were purified by centrifugation and 5% of the lysate was retained as input. The remaining lysate were split into two and its volume were topped up to 100 μL using PXL buffer. 100 μL of protein A Dynabeads (Thermo Fisher) primed with either anti-SFPQ antibody (ab38148) or anti-IgG antibody (MA5-14453) were added to each lysate and incubated at 4°C for an hour on a rotator. Bound SFPQ-RNA complexes were purified and washed thrice in high salt wash buffer (50 mM Tris-HCL, pH 7.4, 1M NaCl, 1mM EDTA, 1% Igepal, 0.1% SDS, 0.5% sodium deoxycholate). Subsequently, bound complexes were washed twice in PNK wash buffer (20 mM Tris-HCL, pH 7.4, 10 mM MgCl_2_, 0.2% Tween-20) and followed by proteinase K digestion (Thermo Fisher). Bound RNAs were purified using phenol-chloroform extraction followed by reverse-transcription to generate cDNAs. Relative amounts of SFPQ-bound RNAs was quantified by qPCR using primers: (Table S9).

### CRISPR/Cas9

gRNAs were formed from chemically synthesized Alt-R®-modified crRNAs from Integrated DNA Technologies (IDT). Each crRNA was suspended in duplex buffer to 100μM concentration, then a crRNA:tracrRNA duplex was formed by combining 3μl crRNA, 3μl 100μM tracrRNA, and 19 μl duplex buffer at 95**°**C for five minutes, then cooled to room temperature and stored at −20**°**C. To make gRNA:Cas9 RNP complexes, a mix was formed as follows: 1.5 μl each gRNA, 0.75 μl 2M KCl, 1.25 μl EnGen Spy Cas9 NLS (NEB). The mix was incubated at 37**°**C for five minutes, then brought to room temperature. One nanoliter of the gRNA:Cas9 complex was injected into embryos at the 1-cell stage. The following gRNAs were used:

*b4galt2*: AAGGATGAATTGAAGGTCAC, AAAGACTTTGTGTGCAACTC
*gdf11*: GTAGAGAGTAGGTTCAGAGT, GACCAAATGTTGTTAGAAAG

### RNA and morpholino injections

The *epha4b* cryptic transcript was amplified from cDNA and inserted into the multi-cloning site of plasmid pCS2+ (Addgene). The *in-vitro* transcription reaction was performed on linearized plasmid using the mMessage mMachine SP6 Transcription Kit (ThermoFisher), and the RNA was purified using a Mini Quick Spin Column (Roche). 100 pg RNA was injected into the embryo at the one-cell stage.

For morpholino knockdown of the *epha4b* cryptic exon, embryos were injected into the yolk at the one-cell stage with 0.1 pmol of Epha4b splice junction morpholino or mismatch.

Epha4b splice junction morpholino: ACAGCTGAGAAAAAAACACGGATAT
Epha4b splice junction mismatch morpholino: ACAcCTcAGAAAtAAAgACcGATAT

### In-situ hybridization

Linearized plasmids containing the antisense sequence for *rfng* (Cheng *et al*, 2004), *deltaA* (Allende & Weinberg, 1994), or the *epha4b* cryptic exon were transcribed into RNA probes using DIG labeling mix (Roche) according to the manufacturer’s instructions. Probes were purified using Mini Quick Spin Columns (Roche). *In-situ* hybridization reaction was performed as described elsewhere (Thomas-Jinu & Houart, 2013).

### qPCR

RNA was extracted from 24-28 hpf *sfpq−/−* embryos and heterozygous or WT siblings using the RNease Mini Kit (Qiagen). 1 ug of extracted RNA was used in a reverse transcriptase reaction using the Superscript III First Strand cDNA Synthesis Kit (Invitrogen). 250 ng of cDNA was used in qPCR reactions with the LightCycler 480 SYBR Green I Master Mix (Roche). Each sample was compared against a B-actin control reaction.

### Bioinformatics

For analyses of 24 hpf *sfpq*^-/-^ RNA-seq data using Cufflinks package (Trapnell *et al*, 2012), reads were mapped to zebrafish GRCz9 assembly and differential expression analysis were carried out using default settings.

For analyses of 24 hpf *sfpq*^-/-^ RNA-seq data using Whippet pipeline (Sterne-Weiler *et al*, 2018), a GRCz10 Ensembl-based index was generated using Whippet’s index building function from the Ensembl-based fasta (ftp://ftp.ensembl.org/pub/release-91/fasta/danio_rerio/dna/Danio_rerio.GRCz10.dna.toplevel.fa.gz) and gene annotation files (ftp://ftp.ensembl.org/pub/release-91/gtf/danio_rerio/Danio_rerio.GRCz10.91.gtf.gz). Quantification of aligned RNA-seq reads were done as follows:

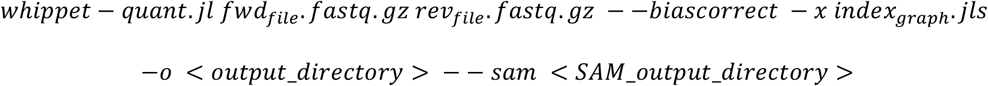

The above quantification function outputs several tables containing read counts at the gene and isoform level. Differential gene and isoform expression analyses were identified using the edgeR package with the estimateGLMRobustDisp function (Robinson *et al*, 2010). Differential splicing events were identified using Whippet’s delta analysis function with default parameters. An event with a “Probability” score exceeding 80% is classified as significantly regulated. Cryptic splicing events were annotated using custom R-scripts.

For analyses of conditional Sfpq knock-out mouse model (Takeuchi *et al*, 2018) dataset, the above Whippet pipeline was carried out using Ensembl’s GRCm38 fasta (ftp://ftp.ensembl.org/pub/release-99/fasta/mus_musculus/dna/Mus_musculus.GRCm38.dna.toplevel.fa.gz) and annotation (ftp://ftp.ensembl.org/pub/release-99/gtf/mus_musculus/Mus_musculus.GRCm38.99.gtf.gz) files. For analyses of conditional ALS-derived iPSC differentiation dataset (Luisier *et al*, 2018), the above Whippet pipeline was carried out using Ensembl’s GRCh37 fasta (ftp://ftp.ensembl.org/pub/grch37/current/fasta/homo_sapiens/dna/Homo_sapiens.GRCh37.dna.toplevel.fa.gz) and annotation (ftp://ftp.ensembl.org/pub/release-75/gtf/homo_sapiens/Homo_sapiens.GRCh37.75.gtf.gz) files.

To construct CLE-containing transcripts, read alignments from Whippet were sorted, indexed and assembled using the StringTie program (Kovaka *et al*, 2019). Ensembl’s GRCz10 transcriptome was used as reference and assembly was done for each biological replicate as follows:

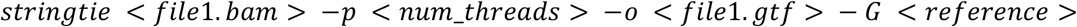

Assembled transcripts from each sample were subsequently combined using StringTie’s merge function using GRCz10 annotations as reference. CLE-containing isoforms were identified by intersecting exon coordinates from the merged transcript assembly with CLE coordinates from Whippet delta analysis output. Intersection operation was done in R using Bioconductor’s GenomicRanges package (Lawrence *et al*, 2013). Analyses on the coding potential of CLE isoforms and its functional loss of protein domains were carried out using custom R-scripts.

For the analyses of introns from which the CLEs were spliced from, intronic features were extracted from the custom-assembled transcript in R using Bioconductor’s GenomicFeatures package (Lawrence *et al*, 2013). A list of the largest, non-overlapping introns was generated and annotated for an overlap with a CLE segment using GenomicRanges’ reduce and subsetByOverlaps functions respectively. The relative position of CLEs within its intron was determined using psetdiff operation followed by extracting the width of the upstream intronic segment.

For the analyses of CLE conservation, 8-way PhastCons data were downloaded from UCSC (http://hgdownload.soe.ucsc.edu/goldenPath/danRer7/phastCons8way/fish.phastCons8way.bw). Coordinates of CLE containing intronswere converted to GRCv9 using UCSC’s LiftOver function and binned into 1 kb sequence using a sliding window technique (1 bp steps). Average PhastCons score of each bin was calculated using bedtools’ “map” function and bins containing CLE were annotated through intersection. Conservation scores of each CLE and 250 nt of its flanking introns were calculated using the same tool. To refine the conservation regions surrounding the intron-CLE borders, average PhastCons score were calculated for 10 nt windows including 30 nt of each exonic ends.

For the analyses of SFPQ binding motifs within sequences surrounding CLEs, its Position-Specific Scoring Matrix was downloaded from RBPmap (http://rbpmap.technion.ac.il/download.html) and manually converted into a MEME motif format (http://meme-suite.org/doc/meme-format.html). Occurrence of SPFQ binding sites was analyzed using MEME’s FIMO program (http://meme-suite.org/doc/fimo.html) using the following parameters:

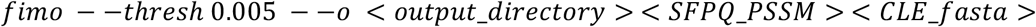

The average number of SFPQ binding motifs were calculated for 25 nt windows of flanking intronic sequence including 25 nt of each exonic ends.

## References

Allende ML & Weinberg ES (1994) The expression pattern of two zebrafish achaete-scute homolog (ash) genes is altered in the embryonic brain of the cyclops mutant. Dev. Biol. 166: 509–530

Blasco H, Bernard-Marissal N, Vourc’h P, Guettard YO, Sunyach C, Augereau O, Khederchah J, Mouzat K, Antar C, Gordon PH, Veyrat-Durebex C, Besson G, Andersen PM, Salachas F, Meininger V, Camu W, Pettmann B, Andres CR & Corcia P (2013) A Rare Motor Neuron Deleterious Missense Mutation in the *DPYSL3* (*CRMP4*) Gene is Associated with ALS. Hum. Mutat. 34: 953–960

Blazquez L, Emmett W, Faraway R, Pineda JMB, Bajew S, Gohr A, Haberman N, Sibley CR, Bradley RK, Irimia M & Ule J (2018) Exon Junction Complex Shapes the Transcriptome by Repressing Recursive Splicing. Mol. Cell 72: 496–509.e9

Bottini S, Hamouda-Tekaya N, Mategot R, Zaragosi LE, Audebert S, Pisano S, Grandjean V, Mauduit C, Benahmed M, Barbry P, Repetto E & Trabucchi M (2017) Post-transcriptional gene silencing mediated by microRNAs is controlled by nucleoplasmic Sfpq. Nat. Commun. 8: 1189

Cagnetta R, Frese CK, Shigeoka T, Krijgsveld J & Holt CE (2018) Rapid Cue-Specific Remodeling of the Nascent Axonal Proteome. Neuron 99: 29–46.e4

Cheng Y-C, Amoyel M, Qiu X, Jiang Y-J, Xu Q & Wilkinson DG (2004) Notch Activation Regulates the Segregation and Differentiation of Rhombomere Boundary Cells in the Zebrafish Hindbrain. Dev. Cell 6: 539–550

Ciolli Mattioli C, Rom A, Franke V, Imami K, Arrey G, Terne M, Woehler A, Akalin A, Ulitsky I & Chekulaeva M (2019) Alternative 3’ UTRs direct localization of functionally diverse protein isoforms in neuronal compartments. Nucleic Acids Res. 47: 2560–2573

Cooke JE, Kemp HA & Moens CB (2005) EphA4 Is Required for Cell Adhesion and Rhombomere-Boundary Formation in the Zebrafish. Curr. Biol. 15: 536–542

Cosker KE, Fenstermacher SJ, Pazyra-Murphy MF, Elliott HL & Segal RA (2016) The RNA-binding protein SFPQ orchestrates an RNA regulon to promote axon viability. Nat. Neurosci. 19: 690–696

Deshaies J-E, Shkreta L, Moszczynski AJ, Sidibé H, Semmler S, Fouillen A, Bennett ER, Bekenstein U, Destroismaisons L, Toutant J, Delmotte Q, Volkening K, Stabile S, Aulas A, Khalfallah Y, Soreq H, Nanci A, Strong MJ, Chabot B & Vande Velde C (2018) TDP-43 regulates the alternative splicing of hnRNP A1 to yield an aggregation-prone variant in amyotrophic lateral sclerosis. Brain 141: 1320–1333

Dye BT & Patton JG (2001) An RNA recognition motif (RRM) is required for the localization of PTB-associated splicing factor (PSF) to subnuclear speckles. Exp. Cell Res. 263: 131–144

Furlanis E, Traunmüller L, Fucile G & Scheiffele P (2019) Landscape of ribosome-engaged transcript isoforms reveals extensive neuronal-cell-class-specific alternative splicing programs. Nat. Neurosci. 22: 1709–1717

Gerety SS & Wilkinson DG (2011) Morpholino artifacts provide pitfalls and reveal a novel role for pro-apoptotic genes in hindbrain boundary development. Dev. Biol. 350: 279–289

Guvenek A & Tian B (2018) Analysis of alternative cleavage and polyadenylation in mature and differentiating neurons using RNA-seq data. Quant. Biol. 6: 253–266

Hall-Pogar T, Liang S, Hague LK & Lutz CS (2007) Specific trans-acting proteins interact with auxiliary RNA polyadenylation elements in the COX-2 3′-UTR. RNA 13: 1103–1115

Hanus C & Schuman EM (2013) Proteostasis in complex dendrites. Nat. Rev. Neurosci. 14: 638–648

Heyd F & Lynch KW (2010) Phosphorylation-dependent regulation of PSF by GSK3 controls CD45 alternative splicing. Mol. Cell 40: 126–137

Van Hoecke A, Schoonaert L, Lemmens R, Timmers M, Staats KA, Laird AS, Peeters E, Philips T, Goris A, Dubois B, Andersen PM, Al-Chalabi A, Thijs V, Turnley AM, van Vught PW, Veldink JH, Hardiman O, Van Den Bosch L, Gonzalez-Perez P, Van Damme P, et al (2012) EPHA4 is a disease modifier of amyotrophic lateral sclerosis in animal models and in humans. Nat. Med. 18: 1418–1422

Holt CE & Schuman EM (2013) The central dogma decentralized: New perspectives on RNA function and local translation in neurons. Neuron 80: 648–657

Iijima Y, Tanaka M, Suzuki S, Hauser D, Tanaka M, Okada C, Ito M, Ayukawa N, Sato Y, Ohtsuka M, Scheiffele P & Iijima T (2019) SAM68-specific splicing is required for proper selection of alternative 3’UTR isoforms in the nervous system. ISCIENCE

Ishigaki S, Fujioka Y, Okada Y, Riku Y, Udagawa T, Honda D, Yokoi S, Endo K, Ikenaka K, Takagi S, Iguchi Y, Sahara N, Takashima A, Okano H, Yoshida M, Warita H, Aoki M, Watanabe H, Okado H, Katsuno M, et al (2017) Altered Tau Isoform Ratio Caused by Loss of FUS and SFPQ Function Leads to FTLD-like Phenotypes. Cell Rep. 18: 1118–1131

Ishigaki S, Masuda A, Fujioka Y, Iguchi Y, Katsuno M, Shibata A, Urano F, Sobue G & Ohno K (2012) Position-dependent FUS-RNA interactions regulate alternative splicing events and transcriptions. Sci. Rep. 2:

Kainov YA & Makeyev E V. (2020) A transcriptome-wide antitermination mechanism sustaining identity of embryonic stem cells. Nat. Commun. 11: 1–18

Ke Y, Dramiga J, Schütz U, Kril JJ, Ittner LM, Schröder H & Götz J (2012) Tau-mediated nuclear depletion and cytoplasmic accumulation of SFPQ in Alzheimer’s and Pick’s disease. PLoS One 7:

Kemp HA, Cooke JE & Moens CB (2009) EphA4 and EfnB2a maintain rhombomere coherence by independently regulating intercalation of progenitor cells in the zebrafish neural keel. Dev. Biol. 327: 313–326

Kim KK, Kim YC, Adelstein RS & Kawamoto S (2011) Fox-3 and PSF interact to activate neural cell-specific alternative splicing. Nucleic Acids Res. 39: 3064–3078

Klim JR, Williams LA, Limone F, Guerra San Juan I, Davis-Dusenbery BN, Mordes DA, Burberry A, Steinbaugh MJ, Gamage KK, Kirchner R, Moccia R, Cassel SH, Chen K, Wainger BJ, Woolf CJ & Eggan K (2019) ALS-implicated protein TDP-43 sustains levels of STMN2, a mediator of motor neuron growth and repair. Nat. Neurosci. 22: 167–179

Knott GJ, Bond CS & Fox AH (2016) The DBHS proteins SFPQ, NONO and PSPC1: A multipurpose molecular scaffold. Nucleic Acids Res. 44: 3989–4004

Kovaka S, Zimin A V., Pertea GM, Razaghi R, Salzberg SL & Pertea M (2019) Transcriptome assembly from long-read RNA-seq alignments with StringTie2. Genome Biol. 20: 278

Langemeier J, Radtke M & Bohne J (2013) U1 snRNP-mediated poly(A) site suppression: Beneficial and deleterious for mRNA fate. RNA Biol. 10: 180–184

Lawrence M, Huber W, Pagès H, Aboyoun P, Carlson M, Gentleman R, Morgan MT & Carey VJ (2013) Software for Computing and Annotating Genomic Ranges. PLoS Comput. Biol. 9: e1003118

Ling JP, Pletnikova O, Troncoso JC & Wong PC (2015) TDP-43 repression of nonconserved cryptic exons is compromised in ALS-FTD. Science (80-.). 349: 650–655

Ling S-C, Polymenidou M & Cleveland DW (2013) Converging Mechanisms in ALS and FTD: Disrupted RNA and Protein Homeostasis. Neuron 79: 416–438

Lowery LA, Rubin J & Sive H (2007) whitesnake/sfpq is required for cell survival and neuronal development in the zebrafish. Dev. Dyn. 236: 1347–1357

Lu J, Shu R & Zhu Y (2018) Dysregulation and Dislocation of SFPQ Disturbed DNA Organization in Alzheimer’s Disease and Frontotemporal Dementia. J. Alzheimer’s Dis. 61: 1311–1321

Luisier R, Tyzack GE, Hall CE, Mitchell JS, Devine H, Taha DM, Malik B, Meyer I, Greensmith L, Newcombe J, Ule J, Luscombe NM & Patani R (2018) Intron retention and nuclear loss of SFPQ are molecular hallmarks of ALS. Nat. Commun. 9: 2010

Martinson HG (2011) An active role for splicing in 3′-end formation. Wiley Interdiscip. Rev. RNA 2: 459–470

Masuda A, Takeda J & Ohno K (2016) FUS-mediated regulation of alternative RNA processing in neurons: insights from global transcriptome analysis. Wiley Interdiscip. Rev. RNA 7: 330–340

Mauger O, Lemoine F & Scheiffele P (2016) Targeted Intron Retention and Excision for Rapid Gene Regulation in Response to Neuronal Activity. Neuron 92: 1266–1278

Melamed Z, López-Erauskin J, Baughn MW, Zhang O, Drenner K, Sun Y, Freyermuth F, McMahon MA, Beccari MS, Artates JW, Ohkubo T, Rodriguez M, Lin N, Wu D, Bennett CF, Rigo F, Da Cruz S, Ravits J, Lagier-Tourenne C & Cleveland DW (2019) Premature polyadenylation-mediated loss of stathmin-2 is a hallmark of TDP-43-dependent neurodegeneration. Nat. Neurosci. 22: 180–190

Mora Gallardo C, Sánchez de Diego A, Gutiérrez Hernández J, Talavera-Gutiérrez A, Fischer T, Martínez-A C & van Wely KHM (2019) Dido3-dependent SFPQ recruitment maintains efficiency in mammalian alternative splicing. Nucleic Acids Res.: 1–14

Nag S, Yu L, Boyle PA, Leurgans SE, Bennett DA & Schneider JA (2018) TDP-43 pathology in anterior temporal pole cortex in aging and Alzheimer’s disease. Acta Neuropathol. Commun. 6: 33

Neumann M, Sampathu DM, Kwong LK, Truax AC, Micsenyi MC, Chou TT, Bruce J, Schuck T, Grossman M, Clark CM, McCluskey LF, Miller BL, Masliah E, Mackenzie IR, Feldman H, Feiden W, Kretzschmar HA, Trojanowski JQ & Lee VM-Y (2006) Ubiquitinated TDP-43 in Frontotemporal Lobar Degeneration and Amyotrophic Lateral Sclerosis. Science (80-.). 314: 130–133

Oh JM, Di C, Venters CC, Guo J, Arai C, So BR, Pinto AM, Zhang Z, Wan L, Younis I & Dreyfuss G (2017) U1 snRNP telescripting regulates a size-function-stratified human genome. Nat. Struct. Mol. Biol. 24: 993–999

Patton JG, Porro EB, Galceran J, Tempst P & Nadal-Ginard B (1993) Cloning and characterization of PSF, a novel pre-mRNA splicing factor. Genes Dev. 7: 393–406

Ray D, Kazan H, Cook KB, Weirauch MT, Najafabadi HS, Li X, Gueroussov S, Albu M, Zheng H, Yang A, Na H, Irimia M, Matzat LH, Dale RK, Smith SA, Yarosh CA, Kelly SM, Nabet B, Mecenas D, Li W, et al (2013) A compendium of RNA-binding motifs for decoding gene regulation. Nature 499: 172–177

Ray P, Kar A, Fushimi K, Havlioglu N, Chen X & Wu JY (2011) PSF suppresses tau exon 10 inclusion by interacting with a stem-loop structure downstream of exon 10. In Journal of Molecular Neuroscience pp 453–466. Humana Press Inc

Robinson MD, McCarthy DJ & Smyth GK (2010) edgeR: a Bioconductor package for differential expression analysis of digital gene expression data. Bioinformatics 26: 139–140

Rosonina E, Ip JYY, Calarco JA, Bakowski MA, Emili A, McCracken S, Tucker P, Ingles CJ & Blencowe BJ (2005) Role for PSF in Mediating Transcriptional Activator-Dependent Stimulation of Pre-mRNA Processing In Vivo. Mol. Cell. Biol. 25: 6734–6746

Sandler JE, Irizarry J, Stepanik V, Dunipace L, Amrhein H & Stathopoulos A (2018) A Developmental Program Truncates Long Transcripts to Temporally Regulate Cell Signaling. Dev. Cell 47: 773–784.e6

Saud K, Cánovas J, Lopez CI, Berndt FA, López E, Maass JC, Barriga A & Kukuljan M (2017) SFPQ associates to LSD1 and regulates the migration of newborn pyramidal neurons in the developing cerebral cortex. Int. J. Dev. Neurosci. 57: 1–11

Shi Y, Di Giammartino DC, Taylor D, Sarkeshik A, Rice WJ, Yates JR, Frank J & Manley JL (2009) Molecular Architecture of the Human Pre-mRNA 3′ Processing Complex. Mol. Cell 33: 365–376

Shi Y & Manley JL (2015) The end of the message: Multiple protein-RNA interactions define the mRNA polyadenylation site. Genes Dev. 29: 889–897

Sibley CR, Emmett W, Blazquez L, Faro A, Haberman N, Briese M, Trabzuni D, Ryten M, Weale ME, Hardy J, Modic M, Curk T, Wilson SW, Plagnol V & Ule J (2015) Recursive splicing in long vertebrate genes. Nature 521: 371–375

Siepel A, Bejerano G, Pedersen JS, Hinrichs AS, Hou M, Rosenbloom K, Clawson H, Spieth J, Hillier LDW, Richards S, Weinstock GM, Wilson RK, Gibbs RA, Kent WJ, Miller W & Haussler D (2005) Evolutionarily conserved elements in vertebrate, insect, worm, and yeast genomes. Genome Res. 15: 1034–1050

Smith A, Robinson V, Patel K & Wilkinson DG (2004) The EphA4 and EphB1 receptor tyrosine kinases and ephrin-B2 ligand regulate targeted migration of branchial neural crest cells. Curr. Biol. 7: 561–570

Sterne-Weiler T, Weatheritt RJ, Best AJ, Ha KCH & Blencowe BJ (2018) Efficient and Accurate Quantitative Profiling of Alternative Splicing Patterns of Any Complexity on a Laptop. Mol. Cell 72: 187–200.e6

Takeuchi A, Iida K, Tsubota T, Hosokawa M, Denawa M, Brown JB, Ninomiya K, Ito M, Kimura H, Abe T, Kiyonari H, Ohno K & Hagiwara M (2018) Loss of Sfpq Causes Long-Gene Transcriptopathy in the Brain. Cell Rep. 23: 1326–1341

Taliaferro JM, Vidaki M, Oliveira R, Olson S, Zhan L, Saxena T, Wang ET, Graveley BR, Gertler FB, Swanson MS & Burge CB (2016) Distal Alternative Last Exons Localize mRNAs to Neural Projections. Mol. Cell 61: 821–833

Thomas-Jinu S, Gordon PM, Fielding T, Taylor R, Smith BN, Snowden V, Blanc E, Vance C, Topp S, Wong CH, Bielen H, Williams KL, McCann EP, Nicholson GA, Pan-Vazquez A, Fox AH, Bond CS, Talbot WS, Blair IP, Shaw CE, et al (2017) Non-nuclear Pool of Splicing Factor SFPQ Regulates Axonal Transcripts Required for Normal Motor Development. Neuron 94: 322–336.e5

Thomas-Jinu S & Houart C (2013) Dynamic expression of neurexophilin1 during zebrafish embryonic development. Gene Expr. Patterns 13: 395–401

Trapnell C, Roberts A, Goff L, Pertea G, Kim D, Kelley DR, Pimentel H, Salzberg SL, Rinn JL & Pachter L (2012) Differential gene and transcript expression analysis of RNA-seq experiments with TopHat and Cufflinks. Nat. Protoc. 7: 562–578

Traunmüller L, Gomez AM, Nguyen TM & Scheiffele P (2016) Control of neuronal synapse specification by a highly dedicated alternative splicing program. Science (80-.). 352: 982–986

Tushev G, Glock C, Heumüller M, Biever A, Jovanovic M & Schuman EM (2018) Alternative 3′ UTRs Modify the Localization, Regulatory Potential, Stability, and Plasticity of mRNAs in Neuronal Compartments. Neuron 98: 495–511.e6

Tyzack GE, Luisier R, Taha DM, Neeves J, Modic M, Mitchell JS, Meyer I, Greensmith L, Newcombe J, Ule J, Luscombe NM & Patani R (2019) Widespread FUS mislocalization is a molecular hallmark of amyotrophic lateral sclerosis. Brain

Venters CC, Oh J-M, Di C, So BR & Dreyfuss G (2019) U1 snRNP Telescripting: Suppression of Premature Transcription Termination in Introns as a New Layer of Gene Regulation. cshperspectives.cshlp.org 11: a032235

Wang G, Yang H, Yan S, Wang C-E, Liu X, Zhao B, Ouyang Z, Yin P, Liu Z, Zhao Y, Liu T, Fan N, Guo L, Li S, Li X-J & Lai L (2015) Cytoplasmic mislocalization of RNA splicing factors and aberrant neuronal gene splicing in TDP-43 transgenic pig brain. Mol. Neurodegener. 10: 42

Wang R, Zheng D, Wei L, Ding Q & Tian B (2019) Regulation of Intronic Polyadenylation by PCF11 Impacts mRNA Expression of Long Genes. Cell Rep. 26: 2766–2778.e6

Wu B, De SK, Kulinich A, Salem AF, Koeppen J, Wang R, Barile E, Wang S, Zhang D, Ethell I & Pellecchia M (2017) Potent and Selective EphA4 Agonists for the Treatment of ALS. Cell Chem. Biol. 24: 293–305

Yarosh CA, Tapescu I, Thompson MG, Qiu J, Mallory MJ, Fu XD & Lynch KW (2015) TRAP150 interacts with the RNA-binding domain of PSF and antagonizes splicing of numerous PSF-target genes in T cells. Nucleic Acids Res. 43: 9006–9016

Zappulo A, Van Den Bruck D, Ciolli Mattioli C, Franke V, Imami K, McShane E, Moreno-Estelles M, Calviello L, Filipchyk A, Peguero-Sanchez E, Müller T, Woehler A, Birchmeier C, Merino E, Rajewsky N, Ohler U, Mazzoni EO, Selbach M, Akalin A & Chekulaeva M (2017) RNA localization is a key determinant of neurite-enriched proteome. Nat. Commun. 8:

