## Supplementary figures for "A conserved role for SFPQ in repression of pathogenic cryptic last exons"

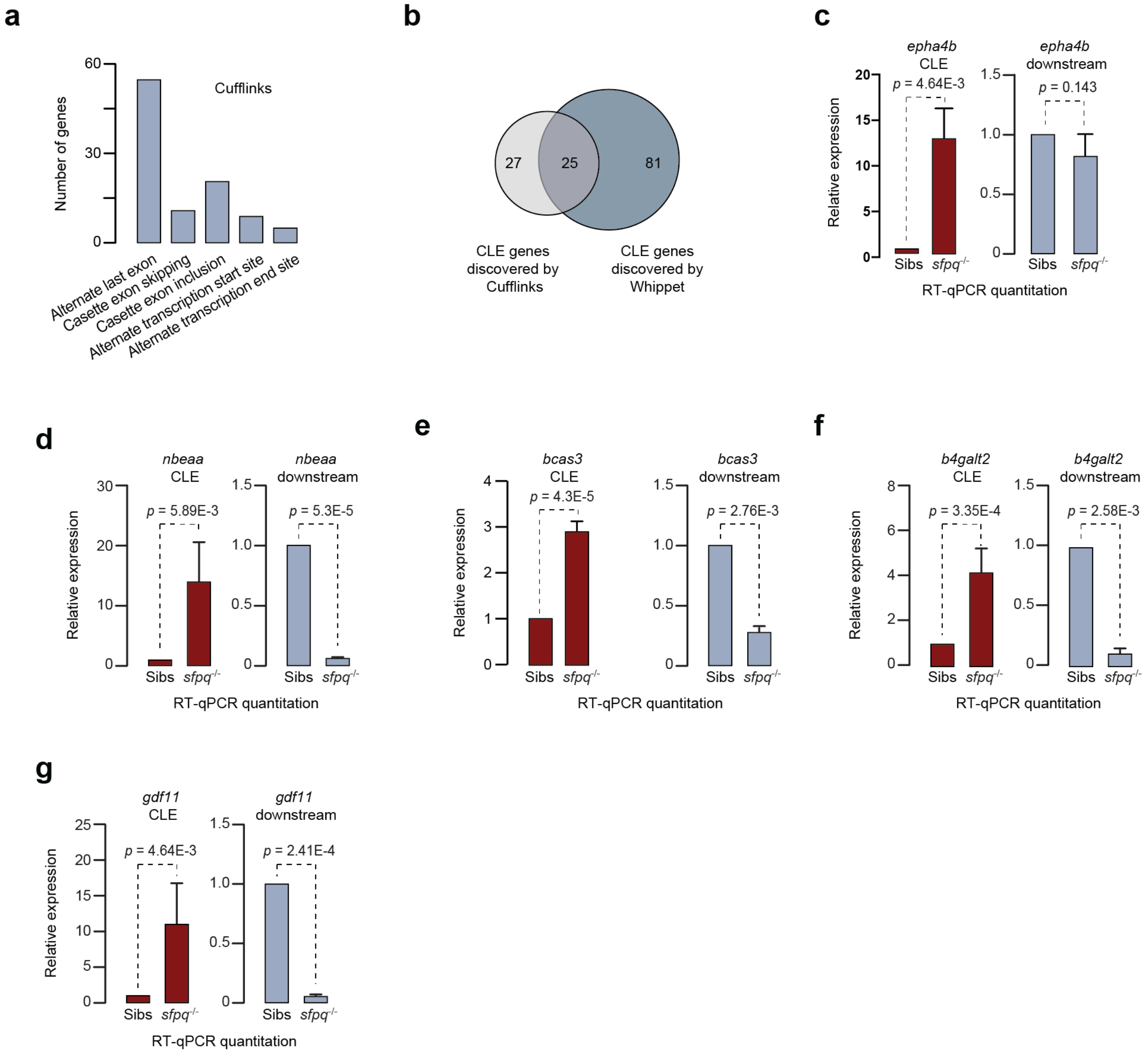


### Fig. S1

**Figure S1: SFPQ regulates the formation of cryptic last exons (CLEs)**
**a**, Bar plot of splicing switches in significantly regulated genes from Cufflinks.
**b**, Total CLE-containing genes found using Cufflinks and Whippet pipelines
**c-g**, RT-qPCR quantitation on the relative expressions of CLE-containing isoforms and CLE-lacking isoforms ("downstream") normalized to β-actin. Expression levels in "Sibs" were set to 1 and mean values from three replicates are shown (± SD).

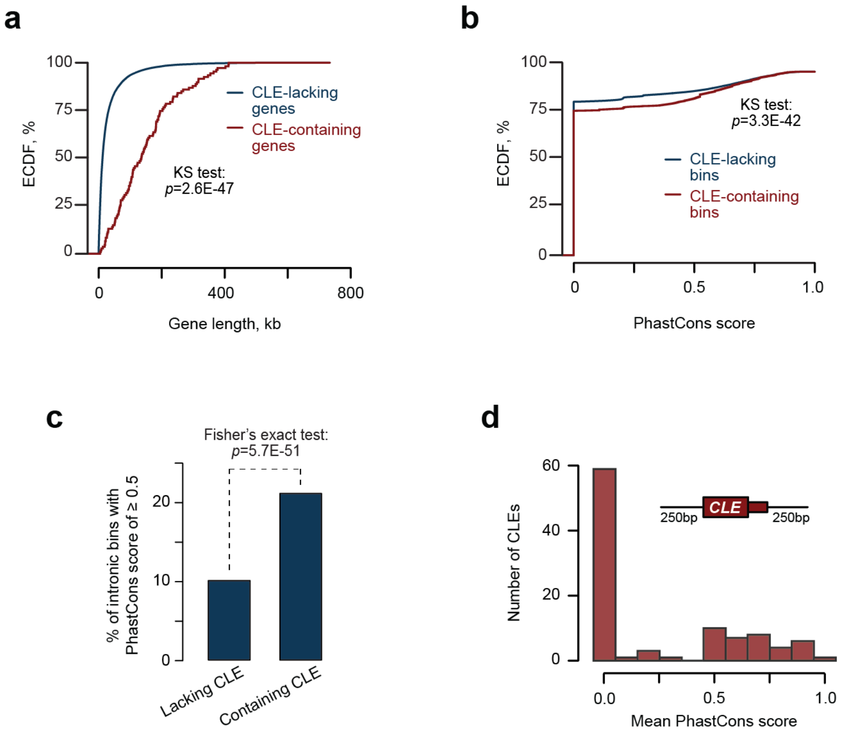


**Fig. S2**

**Figure S2: Properties of CLEs**

**a**, ECDF plot of gene lengths.

**b**, ECDF plot of mean PhastCons scores of bins (1 kb sliding window) along CLE-containing introns.

**c**, Bar plot of the fraction of intronic bins with ≥ 0.5 PhastCons score

**d**, Histogram of mean PhastCons scores of CLE and 250 bp neighbouring its introns.

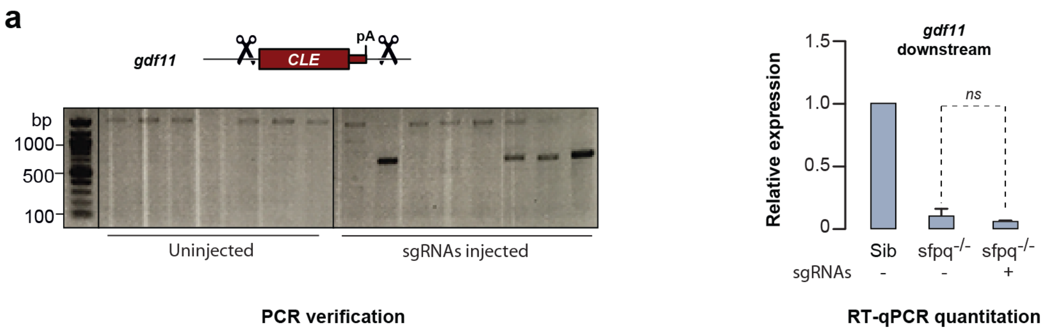


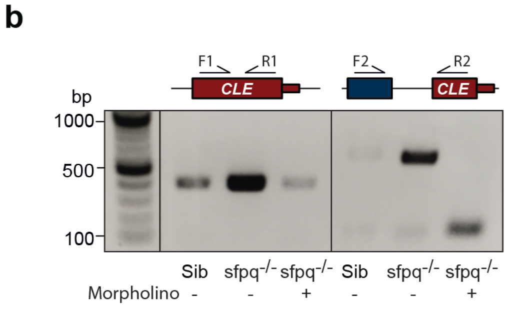


### Fig. S4

**Figure S4: Functional effects of CLEs in sfpq-/- embryos**

**a**, Deletion of the gdf11 CLE using CRISPR/Cas9 does not affect expression levels of the longer isoform. Left: PCR verification of Cas9 cleavage after injection of the gdf11 sgRNAs. Right: RT-qPCR quantitation of the relative expression of the downstream gdf11 exons in sfpq-/- embryos compared to siblings.

**b**, The epha4b splice-junction morpholino prevents expression of the cryptic exon in sfpq-/- embryos

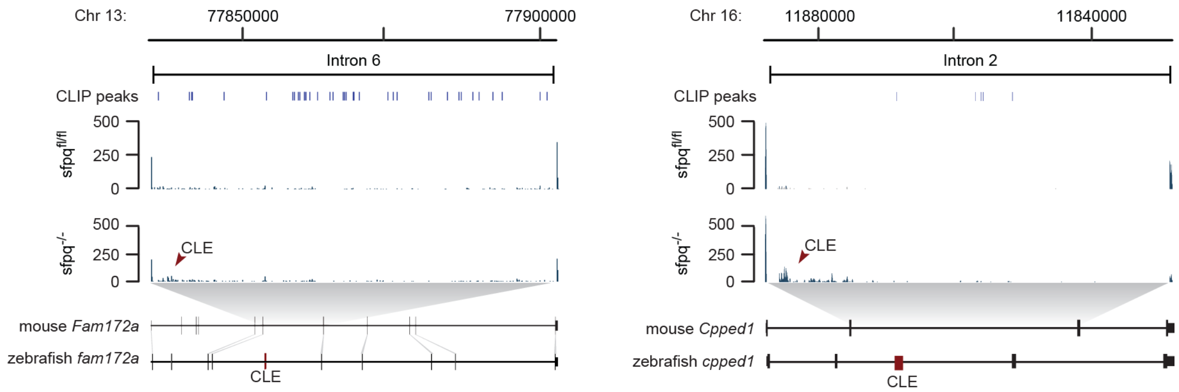


**Fig. S5**

**Figure S5: The CLE-repressing function of SFPQ is conserved in mouse**
Meta-analysis of RNA-seq and CLIP-seq dataset from conditional Sfpq knockout mice for cryptic last exons. Top: distribution of Sfpq CLIP peaks within the CLE-containing intron. Middle: tracks showing read coverage plots and "sashimi" plots from Sfpqfl/fl and Sfpq-/- mice. Bottom: exon architecture of orthologous CLE-expressing genes. Homologous regions between orthologues are shown as connecting lines.
